# The dyslexia susceptibility KIAA0319 gene shows a highly specific expression pattern during zebrafish development supporting a role beyond neuronal migration

**DOI:** 10.1101/267617

**Authors:** Monika Gostic, Angela Martinelli, Carl Tucker, Zhengyi Yang, Federico Gasparoli, Jade-Yi Ewart, Kishan Dholakia, Keith Sillar, Javier Tello, Silvia Paracchini

**Affiliations:** School of Medicine, University of St Andrews, St Andrews, KY16 9TF, UK; Biomedical Sciences Research Complex, University of St Andrews, North Haugh St Andrews, KY16 9ST, UK; College of Medicine and Veterinary Medicine, The University of Edinburgh, Edinburgh EH16 4TJ, UK; SUPA, School of Physics and Astronomy, University of St Andrews, St Andrews, KY16 9SS, UK; School of Psychology and Neuroscience, University of St Andrews, St Andrews, KY16 9JP, UK

**Keywords:** dyslexia, development, zebrafish, gene expression, sensory organs, notochord

## Abstract

Dyslexia is a common neurodevelopmental disorder that affects reading abilities and is predicted to be caused by a significant genetic component. Very few genetic susceptibility factors have been identified so far and amongst those, *KIAA0319* is a key candidate. *KIAA0319* is highly expressed during brain development but its function remains poorly understood. Initial RNA-interference studies in rats suggested a role in neuronal migration whereas subsequent work with double knock-out mouse models for both *Kiaa0319* and its paralogue *Kiaa0319-like* reported effects in the auditory system but not in neuronal migration. To further understand the role of *KIAA0319* during neurodevelopment, we carried out an expression study of the zebrafish orthologue at different embryonic stages. We report particularly high gene expression during the first few hours of development. At later stages, expression becomes localised in well-defined structures such as the eyes, the telencephalon and the notochord, supporting a role for *kiaa0319* that is not restricted to brain development. Surprisingly, *kiaa0319-like*, which generally shows a similar expression pattern, was not expressed in the notochord suggesting a role specific to *kiaa0319* in this structure. This study contributes to our understanding of *KIAA0319* function during embryonic development which might involve additional roles in the visual system and in the notochord. Such a specific spatiotemporal expression pattern is likely to be under the controlled of tightly regulated sequences. Therefore, these data provide a framework to interpret the effects of the dyslexia-associated genetic variants that reside in *KIAA0319* non-coding regulatory regions.

## Introduction

Developmental dyslexia is a specific impairment in learning to read in the absence of any other obvious impairing factors. It affects at least 5% of school-aged children and its heritability is estimated to be above 60% (1). Studying the genetic contribution to dyslexia may help to dissect the underlying neuropsychological mechanisms, which remain hotly debated (2). While a phonological deficit is the most commonly accepted cause for dyslexia, sensory dysfunction in the visual and auditory systems have also been observed in a number of studies (1–3).

The *DYX1C1, DCDC2, ROBO1* and *KIAA0319* genes are known as the classical dyslexia susceptibility genes and they are supported by a number of independent replication studies (4, 5). A role in cortical development, and specifically in neuronal migration, has been proposed for these genes (6), in line with earlier *post-mortem* observations that reported subtle cortical defects in individuals with dyslexia (7, 8). In particular, *KIAA0319* variants have been found to be associated with dyslexia and reading abilities in multiple clinical and epidemiological cohorts (4, 9). A specific dyslexia-associated variant was shown to affect *KIAA0319* transcription regulation and gene expression levels, providing a mechanism to link genetic variation with the disorder (10, 11). Its paralogous gene, *KIAA0319-LIKE* or *KIAA0319L*, has also been reported to be associated with dyslexia (12). Both *KIAA0319* and *KIAA0319L* are transmembrane proteins (13), but their exact cellular functions remain unclear.

In parallel, a new role in cilia biology is emerging for dyslexia candidate genes (14, 15). A transcriptome study showed differential regulation for *KIAA0319, DCDC2* and *DYX1C1* in ciliated tissues (16). Functional studies for Dyx1c1 and Dcdc2 showed a role in ciliogenesis in different biological models (17–19) and some patients with ciliopathies have been found to harbour mutations in both genes (17, 20). While there is no direct evidence supporting a role for KIAA0319 in cilia, the presence of five PKD domains in KIAA0319 lends support to this notion (21). Mutations in PKD genes, which play key roles in cilia, lead to ciliopathies and laterality defects (22). *KIAA0319* has been shown to be a target of T-Brain-1 (TBR1), a transcription factor implicated in autism which regulates different brain developmental processes, such as neuronal migration, axon guidance (23) and the determination of left-right asymmetries in bilaterians (24). *KIAA0319* has been shown to be involved in axon growth and regeneration supporting a role in the adult peripheral nervous system (25).

Both *KIAA0319* and *KIAA0319L* have been implicated in neuronal migration following knockdown experiments that specifically targeted neurons at the early stages of brain development using *in utero* shRNA in rats (11, 26–29). However, knockout (KO) mouse models do not display any cortical abnormalities that could be explained by defective neuronal migration (30, 31). Instead, the KO mice presented auditory system defects (31) in line with observations reported in adult rats that underwent KIAA0319 knock-down *in utero* (27, 32, 33). Therefore, while a role for *KIAA0319* in neurodevelopment is supported by different lines of evidence, its exact function remains largely unknown.

Here, we report a gene expression study for the *kiaa0319* gene during zebrafish development. We observed a specific spatiotemporal expression pattern at specific structures including the eyes and the notochord. Surprisingly, *kiaa0319-like*, which presents a widespread expression in other species, was not expressed in the notochord, suggesting a specific role to *kiaa0319*. These data are suggestive of wider roles for *KIAA0319* during development rather than a specific function restricted to neuronal migration. In particular, our study supports a role outside the brain, specifically in the eyes and in the notochord.

## Materials and Methods

*Fish care*. Wild type zebrafish *(Danio rerio)* (WIK and AB/TU) and the double transgenic Tg(gfap:GFP);Tg(Oligo2:dsRed) were raised at The Queen's Medical Research Institute at the University of Edinburgh according to standard procedures in a Home Office approved facility. Developmental stages, maintained at 28.5°C, were identified as previously described (34). Animals were handled in accordance with the guidelines from European Directive 2010/63/EU and euthanised in accordance with Schedule 1 procedures of the Home Office Animals (Scientific Procedures) Act 1986. Zebrafish embryos were obtained using the Mass Embryo production system (MEPs) of the wild type line Wik. *PCR and qPCR*. Total RNA from developmental stages between 16 – 32 cells, up to 120 hours post-fertilisation (hpf) was extracted using the RNeasy Mini kit according to the manufacturer’s instructions (QIAGEN) using at least 50 embryos at each stage. The heart, liver and brain were dissected from five adult fish, flash-frozen on dry ice and stored at −80°C until the RNA was extracted. Eyes have been dissected and flash frozen on dry ice pooled from 20 individuals (N= 40 eyes total) at 48hpf stage and from five individuals (N = 10 eyes) for the adults.

The PrimeScript RT reagent kit (Takara) was used to transcribe the RNA into the cDNA following the manufacturer’s protocol. The presence of *kiaa0319* transcripts was verified by electrophoresis following PCR amplification. Gene expression level was assessed by quantitative PCR (qPCR) conducted with the Luna Universal RT-qPCR Kit (NEB) and using a Viia7 instrument (Life Technologies, Paisley, UK). For protocol details see Supplementary materials. Primer sequences and accession numbers are shown in Supplementary Table S1.

*Whole-mount in situ hybridization (WISH)*. WISH was carried out following a previously described protocol (35). Briefly, a DIG-labelled riboprobe targeting *kiaa0319* was transcribed using a T3 Polymerase with a DIG RNA Labelling Mix (Roche) according to the manufacturer's instructions and, as a template, a 1066 bp PCR fragment amplified from cDNA (Supplementary Table S1). Zebrafish embryos were collected and processed at 3 somite, 14 somite, 30 hpf and 48 hpf. Embryos were imaged with a Leica MZ16F or MZFLIII bright field microscopes following treatment with an anti-DIG antibody (Roche, diluted 1:5000 in blocking buffer) and a staining solution (NBT and BCIP, Roche).

*RNAscope*. RNAscope ISH (Advanced Cell Diagnostics) was modified from a previously described protocol (36). Samples were fixed in 4% PFA at room temperature for a length of time dependent on the developmental stage (Supplementary Table S2). Samples were hybridized with RNAscope target probes *(kiaa0319l*, nt 545-1425 of ENSDART00000051723.5, Channel 1; *myoD1*, nt 2-1083 of ENSDART00000027661.7, Channel 2; *kiaa0319*, nt 239-1147 of ENSDART00000160645, Channel 3) overnight at 40°C. Images were taken with a Leica TCS SP8 confocal microscope and processed in Leica Application Software X (LAS X) under 20x magnification, unless otherwise specified. Light sheet microscopy (LSM) was conducted with a bespoke microscope built in-house (Supplementary Materials).

## Results

As a first step of our analysis, we verified the expression of *kiaa0319* during early zebrafish development (Supplementary Figure S1). The expression of *Kiaa0319* was observed across all the developmental stages that were analysed. In the adult, expression was much higher in the brain compared to heart and liver, where it is barely detectable. The brain-specific expression pattern is consistent with the expression profile observed for KIAA0319 in humans (Supplementary Figure S2A). In contrast, *KIAA0319L* expression is widespread across human tissues (Supplementary Figure S2B). For a more accurate assessment of expression levels, we analysed *kiaa0319* expression by qPCR and compared it to *kiaa0319l* (Figure 1). Expression of *kiaa0319* was detected across all developmental stages we tested with a similar pattern to *kiaa0319l*. The highest expression was observed for the earliest stages of development (up to 5 hpf) for both genes.

**Figure 1.**
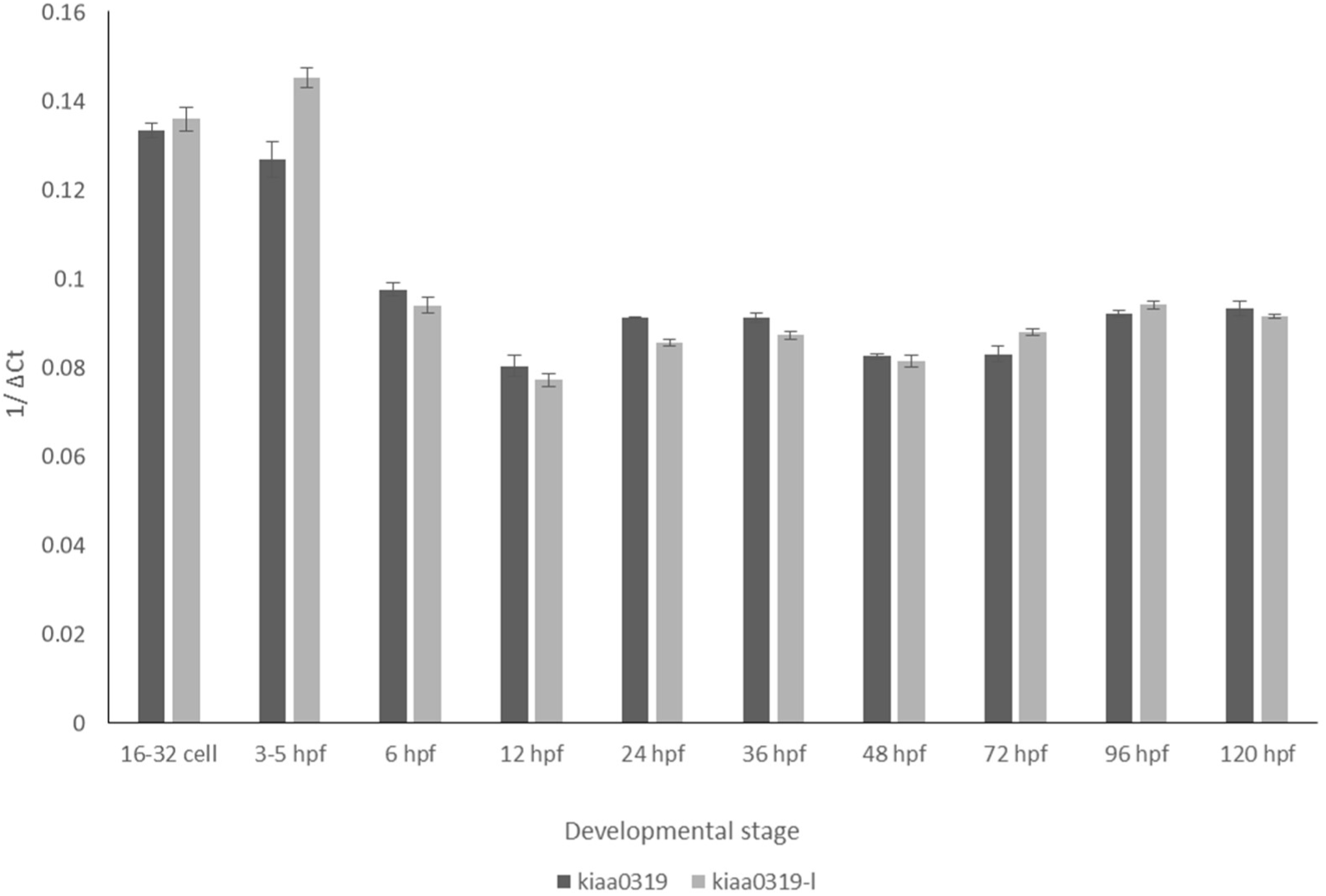
*kiaa0319* is expressed during different stages of development. Quantification of the expression of *kiaa0319* (dark grey) compared to *kiaa0319-like* (light grey) by qPCR measured during the first five days of development. Expression is measured against the *eef1a1l2* gene, used as reference. Results are derived from biological triplicates and error bars indicate standard deviations.

WISH was used to localise the patterns of expression of *kiaa0319* in zebrafish (Figure 2). Consistent with the qPCR data, WISH results show that *kiaa0319* is highly and widely expressed during the early stages of embryonic development (Figure 2A). As development progresses, this widespread expression becomes restricted to specific structures. At the 14 somite stage (16 hpf), *kiaa0319* expression can be visualized in the developing brain and the body midline (Figure 2A.3 and 2A.4). At 30 hpf, expression is detected in the eye, the otic vesicle and in the midbrain-hindbrain boundary (Figure 2B). The midline expression appears localised to the notochord rather than the spinal cord. At 48 hpf expression becomes weaker in the eyes and otic vesicles and is more pronounced in the telencephalon (Figure 2C).

**Figure 2.**
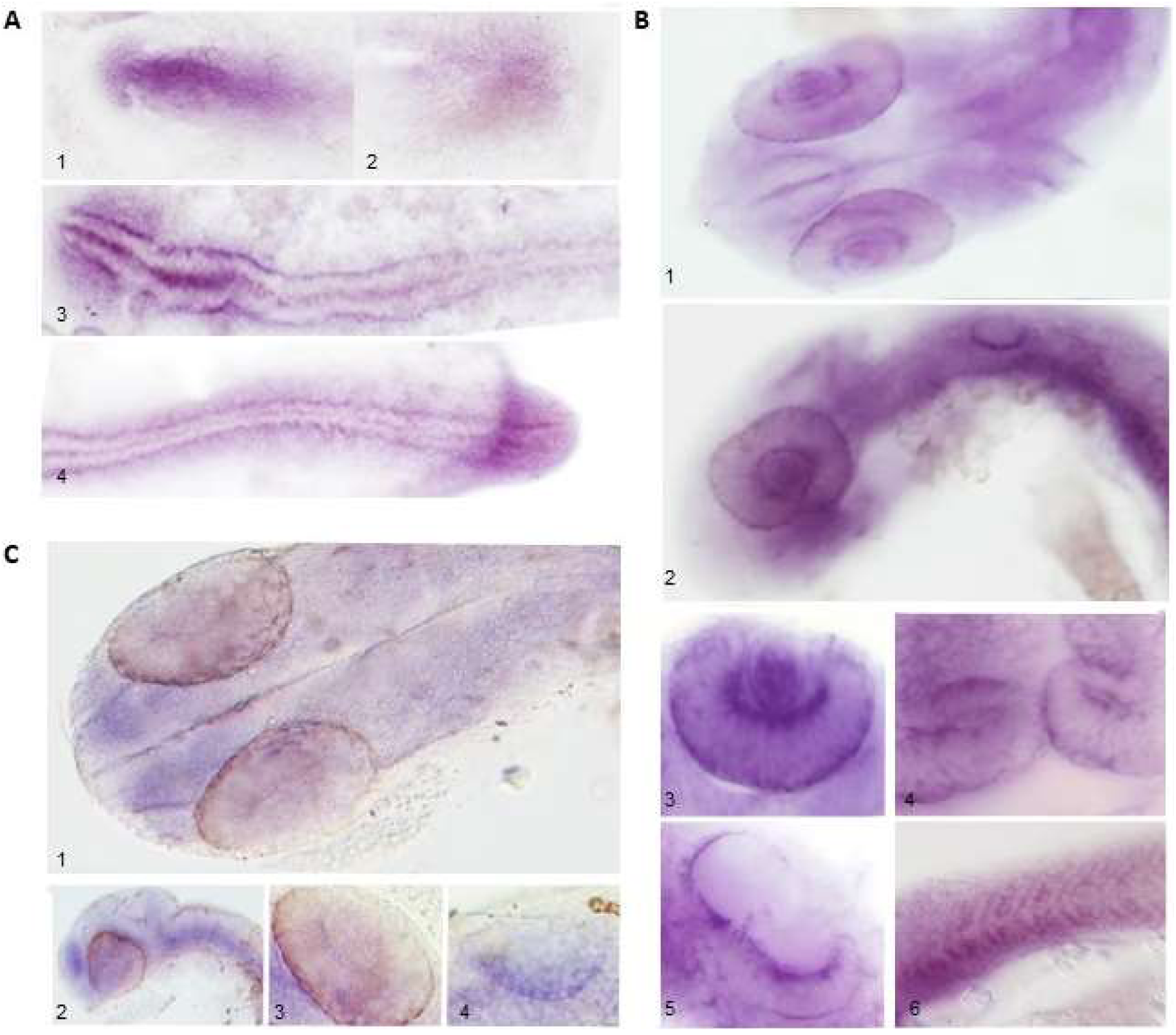
*kiaa0319* presents a specific spatiotemporal expression pattern. **A)** At the 3 somite stage, *kiaa0319* is expressed throughout the embryo with the strongest signal in the head (A1) and to a lesser extent, in the tailbud (A2). At the 14 the somite stage, high expression continues in the head (A3) and is visible along the developing body midline and in the tail (A3 and A4). All images are dorsal views with the head on the left side. **B)** At 30 hpf *kiaa0319* is still expressed throughout the embryo but strong expression emerges in specific structures observed from dorsal (B1) and lateral (B2) views. Details of expression are shown for the eyes (B3; dorsolateral view), the midbrain-hindbrain boundary (B4; dorsal view), the otic vesicles (B5; dorsolateral view) and the notochord (B6; lateral view). **C)** In a dorsal view at 48 hpf, *kiaa0319* expression in the eyes is diminished, while signal in the telencephalon emerges (C1). The signal in the telencephalon is particularly visible in a lateral view (C2) along with expression in the eyes and the region around the notochord. Detailed dorsolateral views of the eye (C3) and otic vesicle (C4) show a specific signal in these structures but at weaker intensity when compared to the pattern observed at 30 hpf.

Both to verify and to achieve higher resolution and specificity we investigated the expression patterns using the highly sensitive RNAscope Fluorescent Multiplex Assay. In particular, we aimed to resolve whether the midline body signal was restricted to the notochord rather than the spinal cord (Figure 3; see Supplementary Figure S3 for the positive and negative controls for these experiments). Consistent with our qPCR and WISH results, at 24 hpf *kiaa0319* expression is widespread but stronger in the brain and body midline. Expression in specific structures, such as the otic vesicles was visible at 48 hpf. RNAscope allowed us to confirm that *kiaa0319* expression was localised to the notochord. For comparison, we analysed the *kiaa0319-like* gene with the same procedure (Figure 3B) and observed a similar expression pattern, with a strong signal in the otic vesicles. However, *kiaa0319-like* gave a much weaker signal in the notochord. Expression of *kiaa0319* in the notochord was further confirmed with fluorescence LSM combined with RNAscope probes targeting *kiaa0319* (Figure 3C). Transverse and lateral images at different developmental stages allowed us to accurately distinguish the spinal cord from the notochord. We detected much higher *kiaa0319* signal intensity in the notochord which became weaker as development progressed. Although weaker, a signal in the spinal cord was also observed. This was strongest at 96 hpf but, rather than increasing or stabilising with development progression, became weaker at 120 hpf. Finally, analysis in the double Tg(gfap:GFP);Tg(Oligo2:dsRed) transgenic embryos further confirmed the *kiaa0319* localization to the notochord. This line is ideal for distinguishing these structures as it presents i) secondary motor neurons, interneurons, and oligodendroglia cells labelled with GFP and ii) motor neurons and oligodendrocytes labelled with DsRed (37).

**Figure 3.**
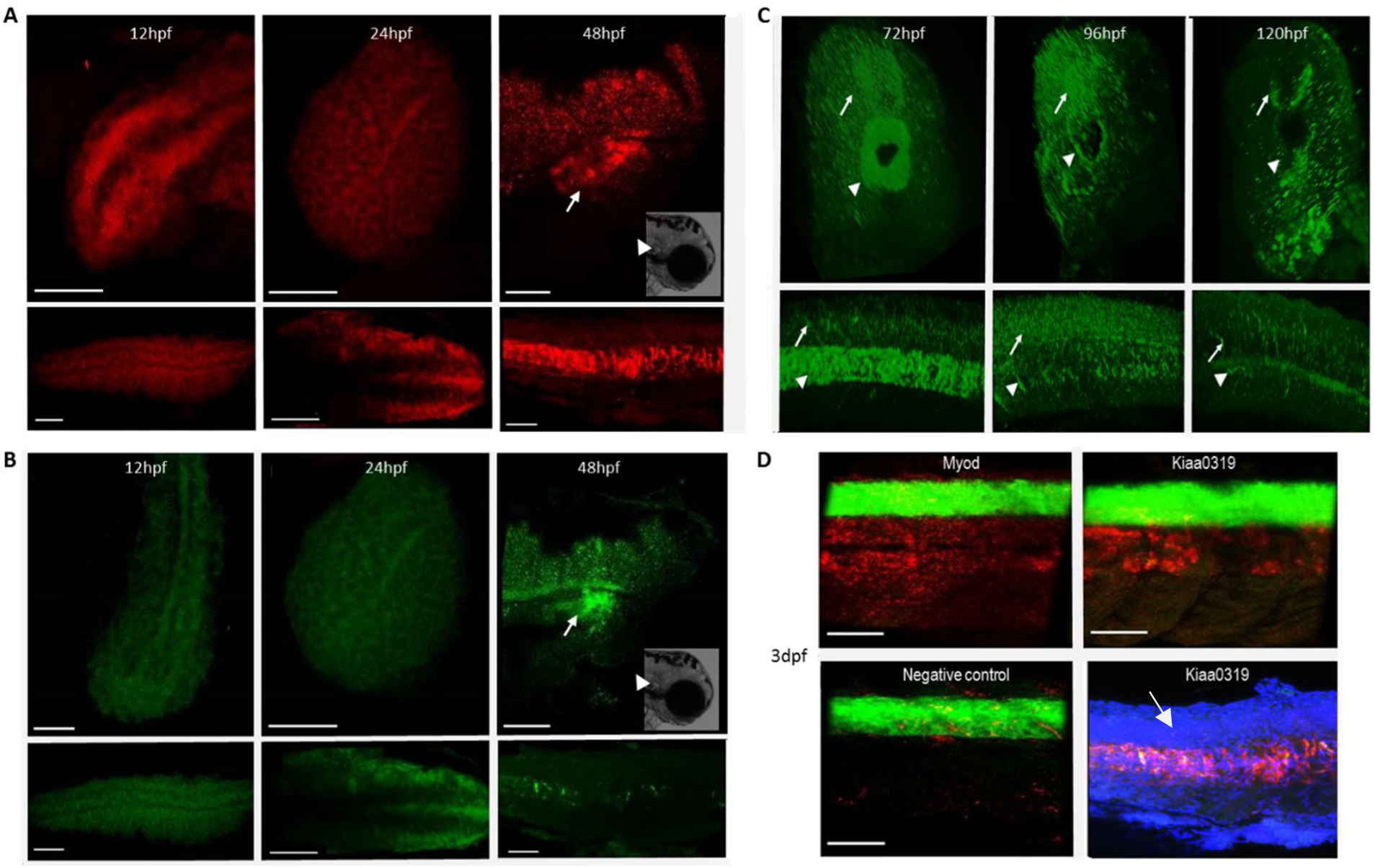
*kiaa0319* is expressed in the notochord, *kiaa0319l* is not. The expression of *kiaa0319* **(A)** and *kiaa0319l* **(B)** was examined at three embryonic stages (12, 24 and 48 hpf) using the RNAScope Fluorescent Multiplex Assay with results shown for the head (top panels) and the body (lower panels). *kiaa0319* (red) is expressed throughout the three stages. High expression is detected in the brain at early hours of development. At 48 hpf, *kiaa0319* is still highly expressed in the brain, in the otic vesicles (top panel; white arrows and white triangle in reference image) and in the notochord (lower panel). *Kiaa0319l* shows a very similar pattern of expression, including a strong signal in the otic vesicles, but the expression in the notochord at 48hpf is much weaker compared to *kiaa0319*. Black area in the brain at 48 hpf correspond to the pigmentation of the embryo. **C)** Transverse (top panels) and lateral (lower panels) 3D reconstructions from light-sheet microscopy images at three developmental stages (72, 96 and 120 hpf) on samples labelled with the *kiaa0319* probe. *Kiaa0319* expression is localised to the notochord (white triangle) but the signal diminishes as development progresses and this transient structure regresses. A well-defined signal in the spinal cord (white arrow) is also detected but it weakens at the later stages of development. **D)** *kiaa0319* shows expression in the notochord. Images collected at 42 hpf by confocal microscopy from the Tg(gfap:GFP);Tg(Oligo2:dsRed) embryos which present GFP (green) in the spinal cord. *myoD1* (myogenic differentiation 1, a universal target for myogenic cells (38)) was used as positive control and demonstrates the specificity of the assay. For reference, the bottom right image shows a wild type embryo treated with the *kiaa0319* RNAscope probe (red) together with DAPI staining (blue). The white arrow indicates the spinal cord where only isolated dots are visible while a strong signal is detected throughout the notochord. The scale bar indicates 50 μm in all panels. For the positive and negative controls see Supplementary Figure S3 and Figure 4.

We also analysed expression in the eyes and at the otic vesicles to investigate whether the signals at these particular structures could have resulted from artefacts such as probe trapping (Figure 4). Both *kiaa0319* and *kiaa0319-like* showed a signal in the eyes (Figure 4A) and at the otic vesicles (Figure 4B), but the latter structure also showed signal in the negative controls. In comparison to the controls, both genes showed stronger expression and a signal characterised by a speckled pattern. While the controls showed a weaker signal, mainly localised along the contour of the otic vesicles, we cannot confirm with confidence that the expression of *kiaa0319* and *kiaa0319l* at these structures. Expression in the eyes was further confirmed by qPCR (Supplementary Figure S4.1).

**Figure 4.**
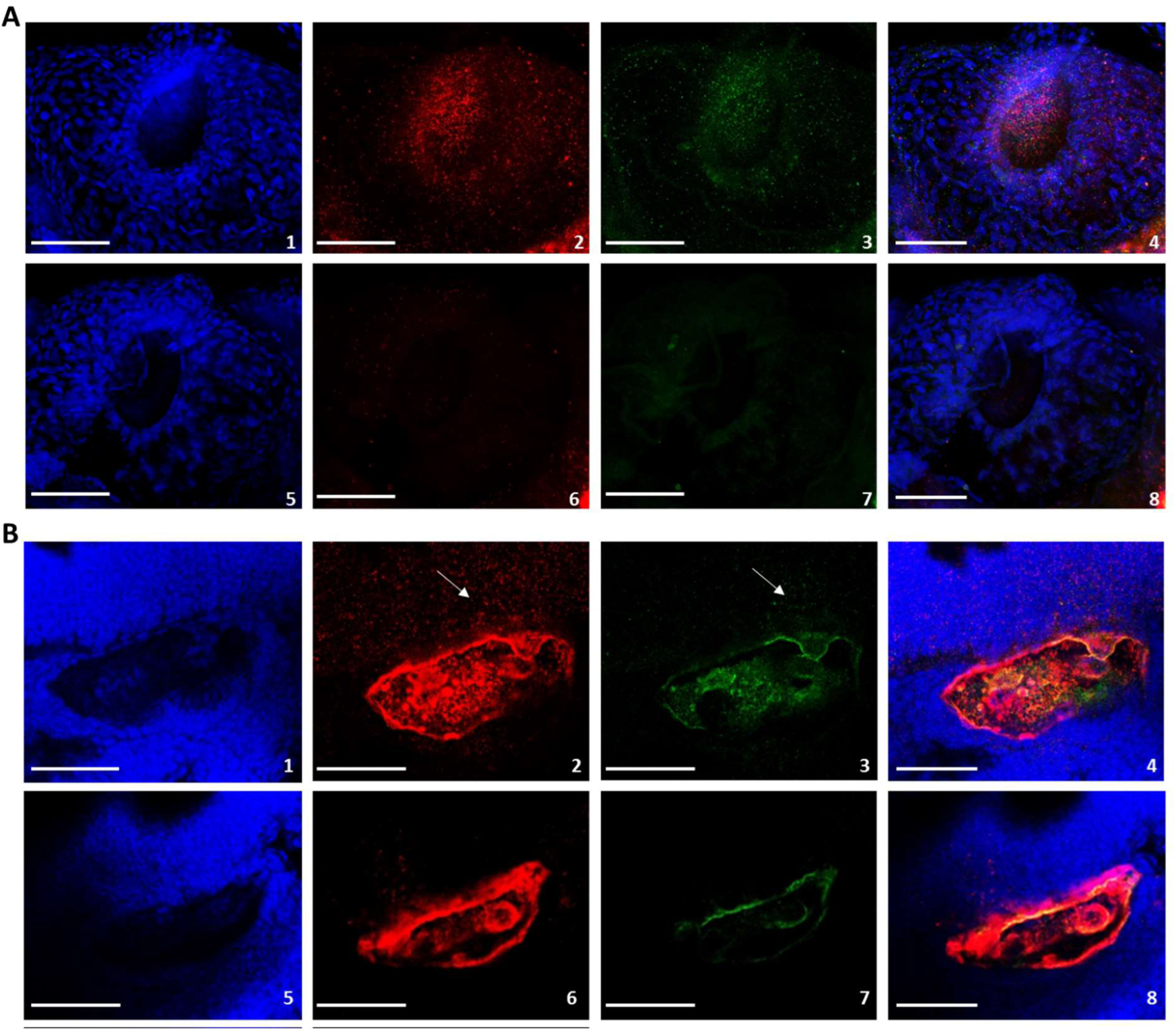
*kiaa0319* and *kiaa0319-like* are expressed in the eyes. RNAscope analysis in the eyes. **(A)** and otic vesicles **(B)** at 48hpf. *kiaa0319*, labelled in red (panels A2 and B2), and *kiaa0319-like*, labelled in green (Alexa 488; panel A3 and B3) are expressed on the surface of the eyes and, most strongly, around the eye lens (see also supplementary Figure S4). The triple negative control shows no signal in the eyes (A6 and A7). Conversely a signal for the triple negative is detected at the otic vesicles (B6 and B7). Expression signal at the otic vesicles appears stronger for both *kiaa0319* (B2) and *kiaa0319-like* (B3) compared to the controls and is detected also in the brain (white arrows; B2 and B3). Expression in the brain is not detected in the negative control, suggesting that probe trapping occurs at the main structure of the otic vesicles. DAPI (first columns; A1, A5, B1 and B5) shows nuclear staining and the last column (A4, A8, B4 and B8) shows the merged signal for all channels. All images show the left side of animals oriented with brain on the left and tail on the right. The scale bar is 50 μm in all panels.

## Discussion

We conducted the first zebrafish characterization of the dyslexia susceptibility *KIAA0319* gene. The expression pattern described in our study supports a specific role for *kiaa0319* in neurodevelopmental processes and adds novel findings towards our understanding of its role in dyslexia. We found that *kiaa0319* is highly expressed at the very first stages of embryonic development with a pattern that becomes increasingly restricted to specific structures such as the telencephalon, the notochord, the eyes and possibly the otic vesicles. For comparison, we analysed also the *kiaa0319-like* gene, which showed a similar pattern but only a very weak signal in the notochord. This observation is surprising given the generally higher and ubiquitous expression of *KIAA0319-LIKE* reported in human tissues (Supplementary Figure S2) and suggests a specific role to kiaa0319. To the best of our knowledge, this is the first study reporting expression of *kiaa0319* during the very first hours of development and clearly showing its expression at specific structures including sensory organs.

Such specific expression pattern is likely to result from a fine-tuned regulation. We showed previously that the *KIAA0319* genetic variants associated with dyslexia reside in the gene regulatory region upstream the promoter (11) and could be pinpointed to a single polymorphism (10). The dyslexia-associated allele of this polymorphism was shown to reduce expression of the *KIAA0319* gene in human cells. It is therefore possible that genetic variation at this locus might also affect the spatiotemporal regulation of expression.

The function of the KIAA0319 protein has been studied in human cell lines and in rodent models, however it is not yet fully understood. The first functional characterization was conducted in rats and suggested a role in neuronal migration (11) while more recent studies in mice indicate an involvement in biological processes beyond brain development (25, 31, 39).

Consistent with the latest studies, we observed expression in the brain, but also observed expression in other organs. Guidi et al (31) generated a double KO mouse model for the *Kiaa0319* and *Kiaa0319l* genes and the most notable phenotype reported was an impairment of the auditory system. Analysis of individual KO for both genes showed mild effects for Kiaa0319l but no effects for Kiaa0319 alone. Rodent models for other dyslexia candidate genes (i.e. *DCDC2* and *DYX1C1)* have also suggested an impairment in auditory processing (40, 41). In this context, the potential expression of both *Kiaa0319* and *Kiaa0319l* in the otic vesicles (Figure 2, Figure 3 and Figure 4) would be in agreement with the data by Guidi and colleagues (31) pointing towards an evolutionarily conserved function. However, further work will be required to establish whether these genes are expressed at the otic vesicles given possible probe trapping in these structures (Figure 4). Given the eye-specific expression observed in our study, it would be expected for vision-related phenotypes to have also been observed in the rodent models. Accordingly, we recommend conducting a thorough visual assessment in future studies of Kiaa0319 knock-out models.

Whether dyslexia is the result of a deficit in sensory systems, as predicted by the magnocellular theory (42), remains highly debated (15). Defects in both the visual and auditory systems have been reported in individuals with dyslexia across different studies, but heterogeneity and inconsistency across studies remain significant challenges (2). The *kiaa0319* expression in the eyes during zebrafish development is in line with a role of sensory organs. While it would be tempting to reach conclusions, it is worth noting that it is not possible to generalise and make strong assumptions based on observations for genes analysed in isolation. Moreover, the *KIAA0319* genetic associations (as with most genetic associations with complex traits) explains only a small fraction of dyslexia heritability (15).

In our study kiaa0319 also shows high expression in the notochord, which instead was very weak for *kiaa0319-like*. The notochord is a transient embryonic structure in zebrafish essential for guiding the development and patterning of the early embryo (43). It is a source of signalling to the surrounding tissues to guide structural development, particularly for the spinal cord. For example, the notochord is the source of sonic hedgehog (SHH) signalling which controls many processes including the development of motor neurons, the establishment of the dorsal-ventral axis and left/right asymmetries (44, 45). The notochord also controls in a highly specific spatiotemporal manner the trajectories of dorsal root ganglion (DRG) axons through repressive signals mediated by aggrecan, one of the chondroitin sulfate proteoglycans (CSPGs) specifically found in the cartilage (46). A similar repressive role has been described for *Kiaa0319* in mice, reporting a role in repressing axon growth in hippocampal and DRG neurons (25). The same study also showed that *Kiaa0319* was expressed in sensory and spinal cord neurons in post-natal and adult mice. Our data are consistent with these findings and support a role for *kiaa0319* both in sensory organs and in body midline structures.

In summary, our characterization of the *KIAA0319* dyslexia susceptibility gene in zebrafish reveals a specific pattern of expression during development. In addition to the expected expression in the brain, we show for the first time high embryonic expression during the first hours of development and, later on, at specific structures including the eyes and in the notochord. These data contribute to our understanding of the role of KIAA0319 during development. Our study supports the emerging role for *KIAA0319* in sensory organs adding to the debate around the different theories aimed at explaining the pathophysiology of developmental dyslexia. In particular, this study will contribute to the ongoing discussion around the role of neuronal migration in dyslexia (39), first proposed in the eighties and corroborated by the initial characterization of susceptibility genes, including *KIAA0319*.

## Acknowledgements

SP is a Royal Society University Research Fellow. This work was supported by Royal Society [RG160373], Carnegie Trust [50341] and Northwood Trust awards to SP. MG was supported by a 600^th^ Anniversary University of St Andrews PhD scholarship and is the recipient of EuFishBioMed scholarship used to visit the Centre de Biologie du Développement (CBD), in Toulouse, France. KD and ZY were supported by a UK Engineering and Physical Sciences Research Council (EPSRC) grant [EP/R004854/1]. The authors are grateful to Prof Caterina Becker for access to zebrafish facilities, to Prof. Bruce Appel for the *Tg(gfap:GFP);Tg(Oligo2:dsRed)* transgenic line, to Drs Patrick Blader and Julie Batut for mentoring on zebrafish experiments during MG stay at the CBD and to Dr Luiz Guidi for comments to the manuscript.

